# Importance of small diameter woody plants and treelet species in tropical wet evergreen forest of southern western ghats, india

**DOI:** 10.1101/2021.12.31.474647

**Authors:** K. A. Sreejith, M.S. Sanil, T.S. Prasad, M. P. Prejith, V.B. Sreekumar, Akhil Murali

## Abstract

Tropical forests have long been accepted for their productivity and ecosystem services on account of their high diversity and stand structural attributes. In spite of their significance, tropical forests, and especially those of Asia, remain understudied. Until recently, most forest inventories in Asia have concentrated on trees 10 cm in diameter. Floristic composition, plant species diversity, above ground biomass, basal area and diversity were investigated across different life form and two-diameter classes in a large scale 10-ha plot, in the undisturbed tropical seasonal rain forest of Southern Western Ghats, Kerala, India. Regeneration pattern of the study area was examined by evaluating fisher’s alpha and IVI (Important Value Index) across three layer of vegetation (seedling, sapling, and tree). Within the plot, we recorded 25,390 woody plant species **≥**1 cm dbh from 45 families, 91 genera, and 106 species. Plant density was 2539 woody individuals per hectare, with a basal area of 47.72 m^2^/ha and an above-ground biomass of 421.77 Mg/ha. By basal area, density, and frequency, the Rubiaceae, Sapotaceae, and Malvaceae families were the most important. Small-diameter trees (1 cm ≤ dbh ≤10 cm) found to be 78 percent of the total tree population, 20.2 percent of the basal area, and 1.4 percent of the aboveground biomass. They also possessed 6 percent more diversity at the family level, 10% more diversity at the genus level, and 12% more diversity at the species level than woody individuals under 10 cm dbh.. Woody individuals of treelets life form and small-diameter classes were much more diverse and dense than the other groups, indicating that results based only on larger canopy trees and larger diameter class may be not an appropriate representation of the diversity status of a particular tropical forest type. Lower density of individuals in initial girth class indicates the vulnerability of the forest system to anthropogenic, natural disturbance and a changing climate. Reduce the minimum diameter limit down to 1 cm, in contrast to 10 cm limit used in most of the evergreen forest inventories, revealed a high density and diversity in thelower stories.

## Introduction

The broad objectives of long-term research are to investigate composition, structure, and dynamics of forest ecosystem on a spatiotemporal scale. Studies from these plots have provided a better understanding of the functioning of the different ecosystems and the knowledge gained is essential for the conservation needs of the ecosystem and strengthening global capacity for forest science research (Condit 1995, Davies et al .2021). Tropical forests are the richest biological communities on earth and these forests have been recognized to harbour a significant proportion of global biodiversity. The consistent positive relationships between stand structural attributes have been reported across moist wet and dry tropical forests (Duran et al. 2015). Stand structural attributes like tree density and tree average diameter had strong positive relationships with above-ground biomass at all spatial scales (Poorter et al. 2015).

Modal plots of majority of plot networks follow 10 cm dbh as the minimum dbh (Blundo et al. 2021). This makes the smaller diameter woody individuals as an overlooked population. Importance of large diameter trees have been studied across various plots (Lutz, J.A. et al. 2021, Bradford and Murphy 2019) The ecological importance of small diameter woody plants to the structure and diversity was determined in tropical evergreen forest at Rabi, Gabon (Memiaghe 2016).

Understory layer is important for the nutrient cycling, biodiversity and regeneration capacity of a tropical forest. Dominant early signals of potential long-term forest responses to climate change are available from studies of understory vegetation there is little information about particularities of diversity, stand structure of understory component of tropical moist forest than higher canopy layer (Claudia and Bryan 2011). In this study, we need to determine the stand structure, diversity and composition of woody plants of different height classes to evaluate the relative contribution of each height class

The Western Ghats is one of the hotspots of biodiversity that support an enormous plant wealth. Results of permanent plots from Western Ghats have been published and comparing to other permanent plots the density of plants in the tropical forest system were reported as moderate and reveals the mature stage of stand and good regeneration (Sukumar et al 1992, Ayyappan and Parthasarathy 1999). Data from large-scale permanent plots of wet evergreen forests of Western Ghats are still lacking which gives an insight on the diversity of species and the distribution of structural features involved in carbon storage and other ecosystem activities. Here our objective is to understand and describe the floristic composition, structure, and physiognomy of a 10 ha. Permanent plot and examine the relative contribution of woody plants to the abundance, basal area, aboveground biomass, and diversity especially in terms of diameter and height class. Regeneration assessment was also done by analyzing the data in different diameter classes.

## Materials and method

### Study site

The study was conducted in the tropical wet evergreen forest at Karadishola which falls in the Sholayar forest range coming under Vazhachal forest division Thrissur, Kerala part of Western Ghats located at, N 10.294485° latitude and, E 76.798551° longitude (Figure. 1). The study site is situated 130 km towards East direction from Chalakkudy in Thrissur District Kerala and 45 km towards west direction from Valparai town, Tamilnadu. The elevation ranges from 900m to 950m above msl. The site receives rain from the southwest (June– September) and the northeast (October–December) monsoons. The vegetation is of tropical wet evergreen forest type. Champion and Seth (1968) classified this under west coast tropical evergreen forest and floristically it is an intermediate type between *Cullenia-Mesua, Palaquium, and Mesua*.Anthropogenic activities in the study area include collection of fuel wood, honey, edible fruits and dammar from *Canarium strictum* and *Vateria indica*.

**Figure 1:**
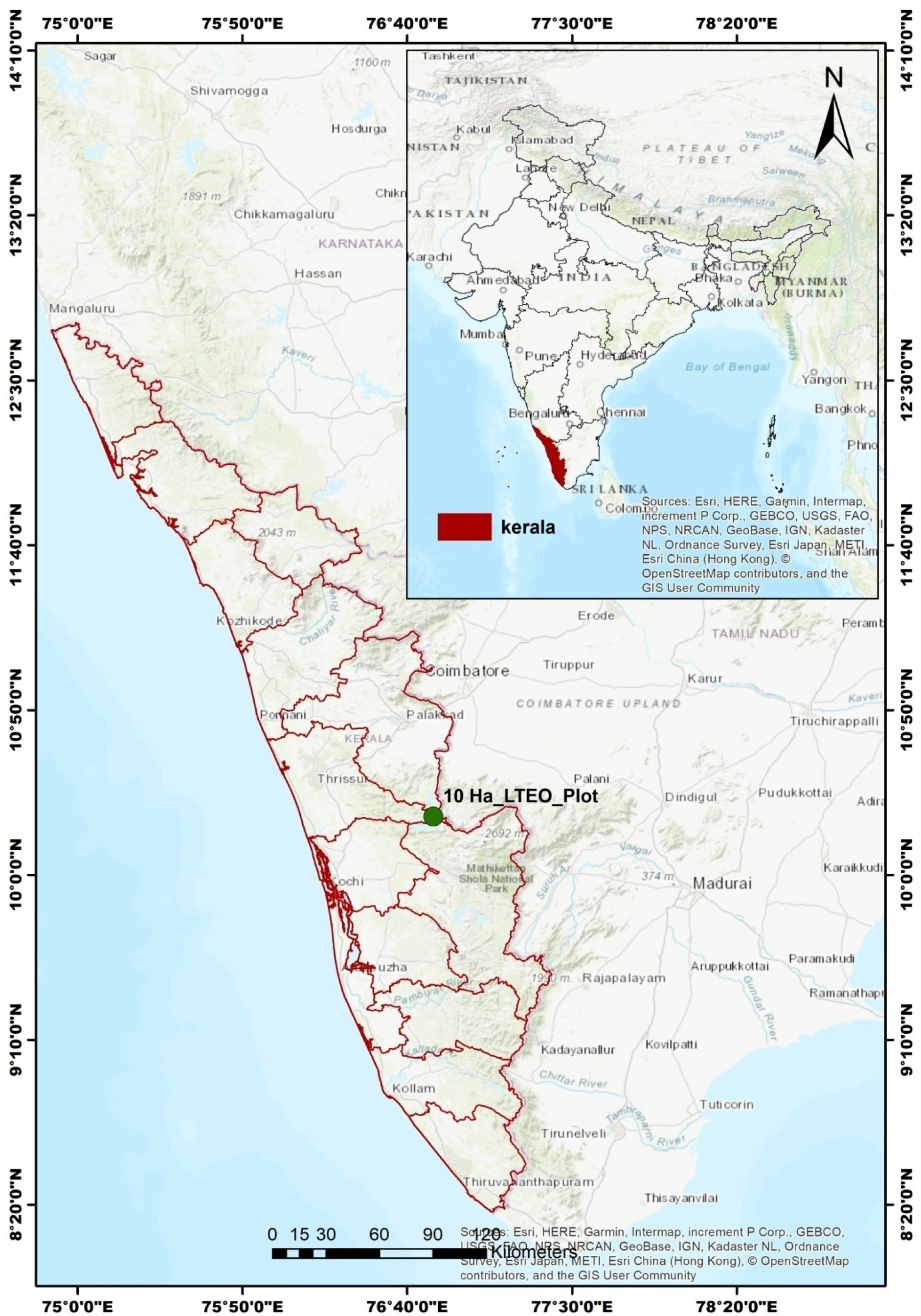
Location map of 10 ha Sholayar LTEO plot in Kerala, India.

## Methodology

### Census

Field survey methods were in accordance with the standards of the Smithsonian Center for Tropical Forest Science Forest-GEO. We first surveyed the 10ha(500×200 m^2^), placing permanent stakes precisely every 20 m with minimal damage to t he vegetation (Condit 1998). Subsequently, all woody individuals ± 1cm diameter-breast-height (dbh) were tagged with numbered aluminum tags, measured, and identified. Stem diameter was measured 1.3 m above the ground, with swollen or buttressed trees measured at a spot where the trunk was more regular; the measurement spot was painted so future measurements could match. Any stem fork or branch <1.3 m above the ground was treated as a secondary stem and measured. Establishment and enumeration (tagging, GBH measurement) of 10 ha plot was carried out from 26 th june 2016 to 28 th December 2018. The individuals were identified using standard floras (Hooker 1872, Gamble and Fischer 1915-1935, Pascal and Ramesh 1987, Sasidharan 2004, Nayar *et al*. 2014) and identified voucher specimens were deposited in the KFRI herbarium.

### Biomass estimation

The aboveground biomass (AGB) of each individual was estimated using the allometric equation proposed by Chave et al. 2014, defined as:

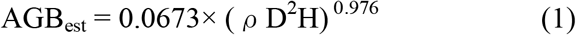

where AGB (kg) is the estimated aboveground biomass, DBH (cm) is the diameter of the tree at breast height, H (m) is the estimated total height, and ρ (g /cm^3^) is the wood density.

To get wood density, we applied the following standardized protocol: (i) Tree identified to the species level were assigned the corresponding wood density value from the Global Wood Density Database (GWDD) (Chave et al. 2009); (ii) When a species was identified at the genus, we used the average density of wood for all species of the same genus in the database and (iii) When trees with no botanical identification or that were not present in the GWDD we used the average density of wood for all species in the plot (Herault and Camille 2018)

Tree height was retrieved by applying the local Height-Diameter (H-D) models; we also used an unpublished dataset of tree height and diameter from an evergreen forest of Southern

Western Ghats. After comparing different local H-D models, we used the method with the lowest residual error to compute the local height–diameter model (Feldpausch et al .2012). Then we retrieved height using this local H-D model. The wood density and height data were retrieved and the aboveground biomass (AGB) was estimated taking all uncertainties into account using the BIOMASS r package (Réjou-Méchain et al. 2017).

### Height categories of species

We classified all species into four growth forms according to their estimated maximum height (Kenfack et al 2007). Treelets and small trees include all species with adult generally less than 10 m tall; understory trees are those with adults 10–20 m tall; lower canopy species have heights 20–30 m; and upper canopy species are those often >30 m in height and emergent above the main canopy. Corresponding adult stem diameters were <10 cm, 10–30 cm, 30–60 cm, and >60 cm dbh, respectively. Information on the heights of the species came from field estimates in the plot supplemented with information from the literature (Sasidharan 2012).

### Data analysis

Because in most of the inventories in the tropical forests are included only individuals with dbh **≥**10 cm, for comparison we report abundance basal area above ground biomass on a per hectare basis as well as for all 10 ha combined and for diameter classes of both 1cm ≤ dbh <10 cm and**≥**10 cm (Memiaghe et al .2016). Standard deviations of each parameter are obtained by dividing the 500×200 m^2^ into 10 non-overlapping 100×100 m^2^ and counting individuals on each. Our counts on individual do not include multiple stems per tree, but multiple stems are included in the basal area calculations..To provide quantitative estimates of plant diversity, fishers alpha, Shannon’s Index Hand Simpson’s index were used. Two-sided, paired t-test was used to test the difference of species diversity indices, biomass, basal area, density between plants with dbh **<10cm** and **≥10cm**. The means of basal area, genera, species and stem per hectare were also calculated for the four life form in each sub plot. One-way analysis of variance (ANOVA) was used to test the differences between the means of these parameters using R package.

### Regeneration assessment

Regeneration status was assessed by evaluating the girth class categories and the categories of trees were based on the girth class of individuals. >30 cm at gbh (girth at breast height) were considered as trees, 10-30 cm gbh as saplings and <10 cm gbh as seedlings(Sushil.2016). The vegetation data were quantitatively analyzed for relative density, relative frequency and relative dominance based on which Important value index(IVI) (Curtis &McIntosh, 1950) was estimated.. Regeneration was characterized with density and dominance. While density is the number of recruits (seedlings and/ or saplings according to the context) in a unit area (SushilSaha .2016). For comparing the diversity species richness and Fisher’s α value were calculated for seedling, sapling and tree. The trees were further divided into seven girth classes i.e. class 30-60, 60-90, 90-120, 120-150, 150-180, 180-210, 210-240 > 240 for density and girth class distribution of trees.

## Results

### Woody floristic structure and diversity

We observed 106 morphospecies inside the 10-ha plot, of which 95 (85.2%) were recognized to species and the remaining 11 (14.8%) to genus level. The 106 morphospecies belongs to 45 families and 91 genera. Euphorbiaceae and Lauraceae were the most diverse family with 8 species in 5 genera, trailed by Meliaceae and Rubiaceae that had 8 species in 4 genera and 7 species in 6 genera respectively.

Anacardiaceae (6 species in 5 genera), Rutaceae (6 species in 5 genera) and Phyllanthaceae (6 species in 4 genera) are the other diverse families in the permanent plot. The genera Aglaia and Mallotus were the richest with 5 and 4 species respectively, followed by Litsea, Aporosa, syzygium each with three species.

Altogether the large-diameter group comprised 86 species in 73 genera and 39 families, while 99 species (96% of the total) in 83 genera and 42 families were recorded among small-diameter trees. In the present study plot 75 species/ha were recorded for all woody individuals, Mean species richness of woody plants was much higher in the small diameter class than that of high diameter class (70 species/ha in class1 and 48 species/ha in cass2) in each plot. Furthermore, the class 1 woody plants are more diverse than the class 2 plants indicated by higher values of other three-diversity index. The shanon diversity index of the class1 was not significantly higher than class2 (P=0.02) (Table 1).

**Figure 1:**
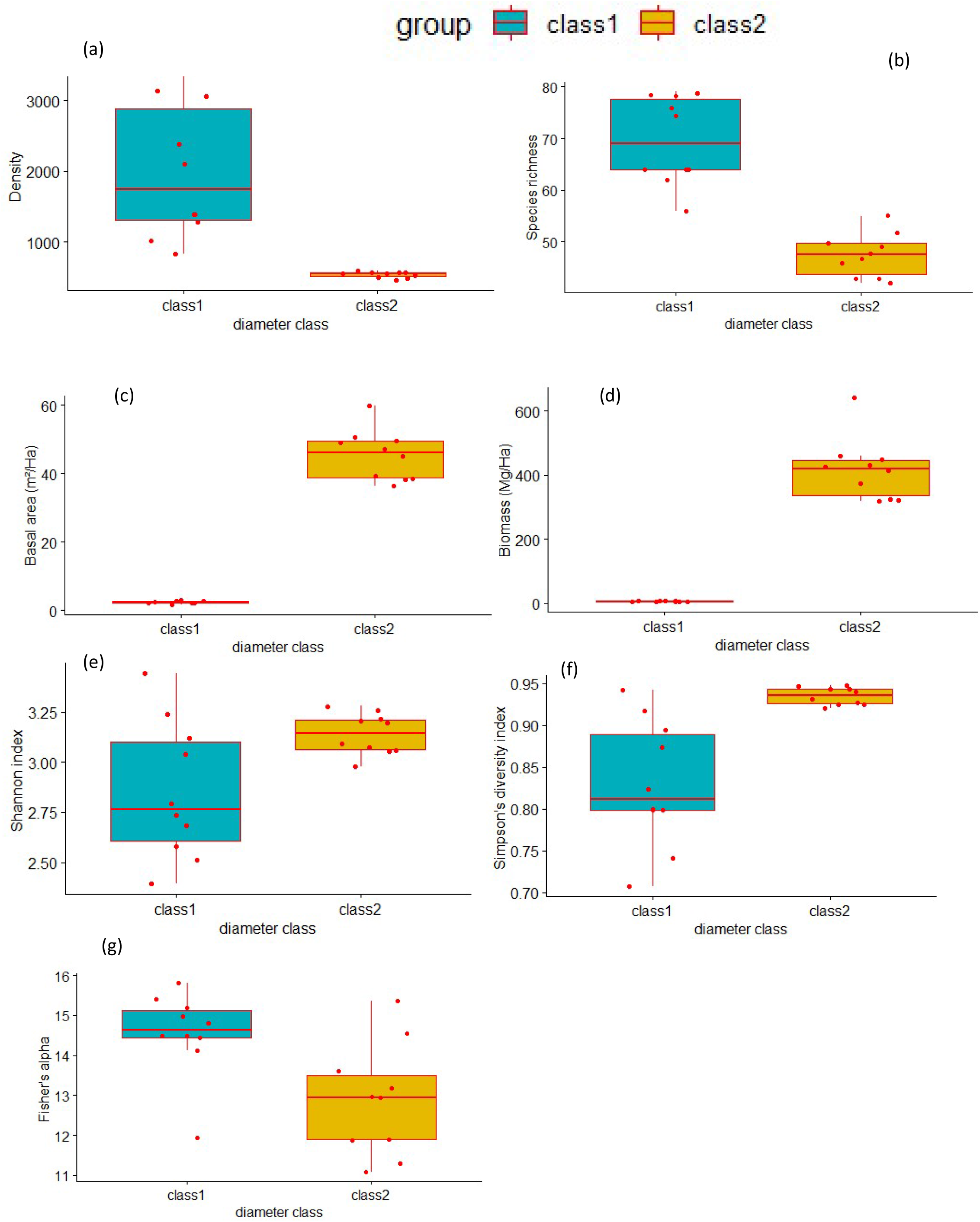
Box plots showing distribution of, density (a), species richness (b), basal area(c), biomass (d), Shannon index (e), Simpson index (f), fisher’s alpha (g) across two diameter classes

**Table 1:** **Comparison of the distribution of structure and diversity across small-diameter (1 cm ≤ dbh <10 cm) and large-diameter (dbh ≥10cm) tree of the sholayar 10-ha plot. Numbers in parenthesis represent standard deviation or percentages.**

| | $\geq 1\text{cm}$ | $< 10\text{cm}$ | $\geq 10\text{cm}$ |
| --- | --- | --- | --- |
| all families | 45 | 42 | 39 |
| all species | 106 | 99 | 86 |
| all genera | 91 | 83 | 73 |
| All trees | 25390 | 19975 | 5415 |
| total basal area | 477.24 | 23.364 | 453.882 |
| total above ground biomass | 4217.779 | 60.847 | 4156.931 |
| mean density per hector | 2539 (975.2) | 1997.4 (946.57) | 541.6 (43.97) |
| mean number of species per hec | 75.3 (8.459) | 69.5 (8.343) | 47.5 (4.196) |
| mean basal area per hec | 47.724 (7.5) | 2.336 (0.42) | 45.38 (7.34) |
| mean AGB per hec | 421.777 (95.93) | 6.0847 (1.037) | 415.693 (95.347) |
| Shannon diversity index | 3.143 | 2.841 | 3.11 |
| Simpson diversity index | 0.123 | 0.171 | 0.066 |
| fisher's alpha index | 15.542 | 14.56 | 12.87 |

### Abundances

In the 10-ha plot, a sum of 25390 trees were recorded, with a moderate density of 2539 individuals/ha. Density of woody species in two diameter class varied significantly (p value < 0.01). Abundance Small-diameter trees was 3.6 times greater than large-diameter trees. There were 19975 small-diameter plants (78.67 % of all stems), averaging 1997 individuals/ha, while large-diameter trees had an abundance of 5415 plants and density of only 546 individuals/ha.. The family Rubiaceae, is the most abundant, with densities >900 individuals/ha, followed by Euphorbiaceae, Urticaceae, Sapotaceae, Meliaceae, Malvaceae, and Putranjivaceae. Among tree genera Palaquium, cullenia, of family Sapotaceae and Malvaceae respectively, were the most abundant, with >100 individuals/ha, followed by Drypetes, Mesua, Aglaia, Vateria, syzygium and Agrostistachys at species level, *Psychotria nudiflora, Palaquium ellipticum, Cullenia exarillata, Vateria indica* were the dominant based on IVI value, followed by *Mesua ferrea and Dendrocnide sinuate*

### Basal Area

The 10 hectare LTEO plot has total basal area of 477.24 m^2^ (47.72± 7.58 m^2^/ha), 20% of the basal area were occupied by Malvaceae (9.69 m2/ha)and Sapotaceae(9.6m^2^/ha) followed by Dipterocarpaceae, Calophyllaceae, Putranjivaceae and Sapindaceae. The dominant genera by basal area were Cullenia, Palaquium, Vateria, Mesua, Drypetes, Dimocarpus. At species level, *Cullenia exarillata* and *Palaquium elipticum* were the most important followed by *Vateria indica, Mesua ferrea, Dimocarpus longan, Drypetes wightii and Holigarna nigra*. Small-diameter trees contributed 4.89% of the total basal area.From the t test, it was found that the difference in the means of basal area among two diameter class were statistically significant at P≤0.01.

### Aboveground biomass

Total aboveground biomass of the 10–ha plot was 4217.779 Mg, averaging 421.777±95.93Mg/ha for all woody plants of **≥**1cm dbh. Malvaceae is reported to be the family with highest AGB (106.97Mg/ha), followed by Sapotaceae (81.7Mg/ha), Calophyllaceae (57.97 Mg/ha), Dipterocarpaceae(47.30781Mg/ha) and Putranjivaceae(23.32Mg/ha). Cullenia and Palaquim were the most important genus in the plot (Table 4). *Cullenia exarillata* was the most important species in terms of aboveground biomass, followed by *Palaquim elipticum, Mesua ferrea* and *Vateria indica*. The aboveground biomass was significantly varied among two-diameter classes at P≤0.01. AGB for large-diameter trees was 415.693±95.347 Mg/ha. whereas the AGB of small diameter class was 6.0847±1.037 Mg/ha, approximately 1.29% of the total biomass (Table 2)

**Table 2:**
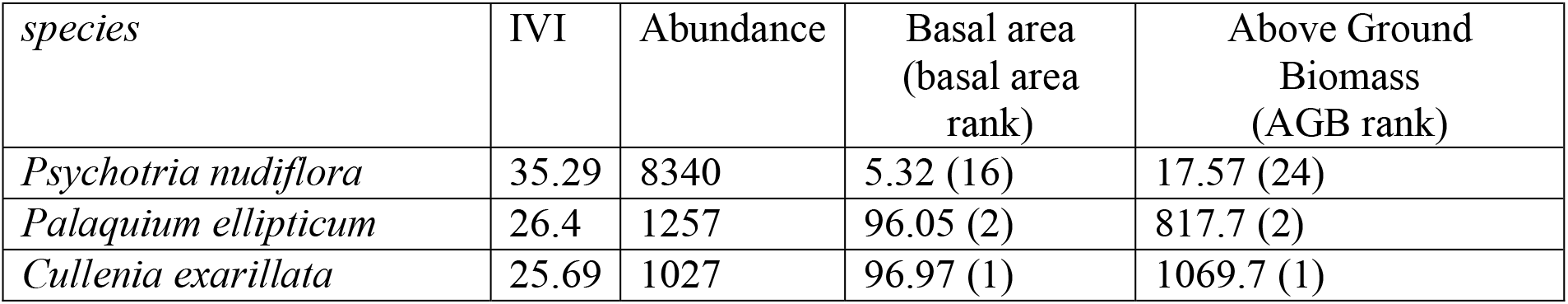
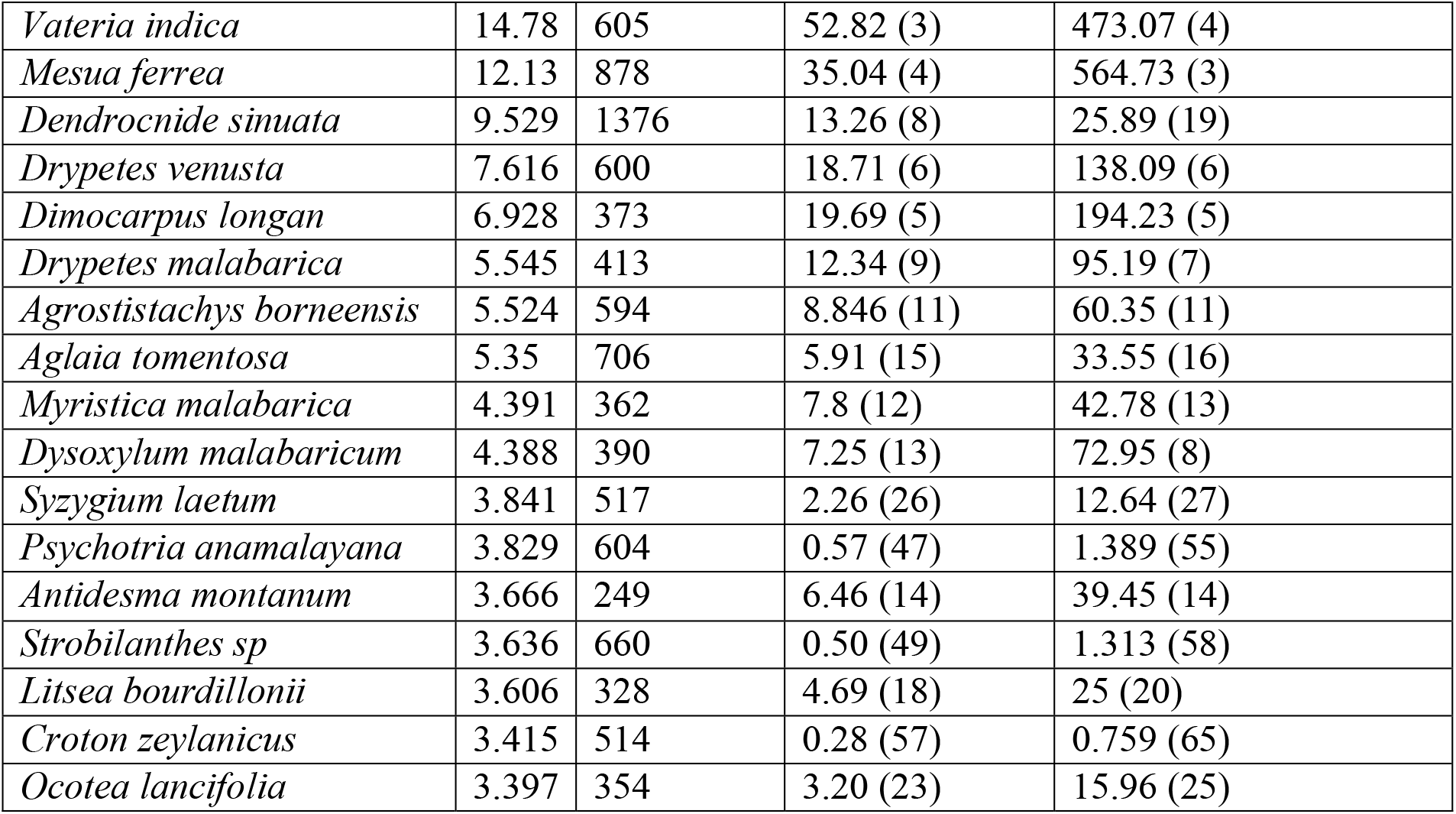
The 20 most dominannt tree species in the 10-ha sholayar plot ranked by IVI. Basal area and AGB rank indicated in parenthesis.

**Table 3:** The 20 most abundant tree genera in the 10-ha sholayar plot ranked by density. Basal area and aboveground biomass (AGB) rank indicated in parenthesis.

| Genus | Abundance | Basal area (m<sup>2</sup>) rank in parenthesis | Biomass (unit) rank in biomass are in paranthesis |
| --- | --- | --- | --- |
| Psychotria | 8944 | 5.88 (18) | 18.96 (23) |
| Dendrocnide | 1376 | 13.26 (8) | 25.89 (19) |
| Palaquium | 1257 | 96.05 (2) | 817.7 (2) |
| Cullenia | 1027 | 96.97 (1) | 1069.7 (1) |
| Drypetes | 1013 | 31.06 (5) | 233.2 (5) |
| Mesua | 878 | 35.04 (4) | 564.7 (3) |
| Aglaia | 845 | 6.50 (15) | 37.11 (15) |
| Strobilanthes | 697 | 0.53 (40) | 1.40 (44) |
| Vateria | 605 | 52.82 (3) | 473. (4) |
| Syzygium | 595 | 3.09 (22) | 20.05 (22) |
| Agrostistachys | 594 | 8.84 (10) | 60.35 (11) |
| Litsea | 550 | 7.01(14) | 35.50 (16) |
| Croton | 514 | 0.28 (46) | 0.759 (51) |
| Aporosa | 404 | 1.12 (30) | 3.509 (39) |
| Lasianthus | 396 | 0.32 (44) | 1.325 (46) |
| Mallotus | 395 | 5.25 (19) | 26 (18) |
| Dysoxylum | 390 | 7.25 (13) | 72.95 (7) |
| Myristica | 390 | 8.72 (11) | 47.5 (13) |
| Meiogyne | 389 | 1.77 (27) | 6.55 (32) |
| Dimocarpus | 373 | 19.69 (6) | 194.23 (6) |

**Table 4:** The 15 most abundant Plant families in the 10-ha Sholayar plot, ranked by FIV (Family Important value index).

| family | species | D | Ba | RD | RDo | RDi | FIV | AGB |
| --- | --- | --- | --- | --- | --- | --- | --- | --- |
| Rubiaceae | 7 | 936.4 | 0.62 | 36.96 | 1.31 | 6.60 | 44.87 | 20.31 (20) |
| Sapotaceae | 2 | 129 | 9.61 | 5.09 | 20.25 | 1.89 | 27.23 | 817.8 (2) |
| Malvaceae | 1 | 102.7 | 9.70 | 4.05 | 20.44 | 0.94 | 25.43 | 1069.7 (1) |
| Euphorbiaceae | 8 | 153.9 | 1.59 | 6.07 | 3.35 | 7.55 | 16.97 | 94.42 (8) |
| Meliaceae | 8 | 124.1 | 1.38 | 4.90 | 2.90 | 7.55 | 15.35 | 110.12 (7) |
| Dipterocarpaceae | 1 | 60.5 | 5.28 | 2.39 | 11.13 | 0.94 | 14.46 | 473.07 (4) |
| Lauraceae | 8 | 97 | 1.13 | 3.83 | 2.38 | 7.55 | 13.76 | 58.46 (12) |
| Calophyllaceae | 2 | 89.3 | 3.71 | 3.52 | 7.81 | 1.89 | 13.22 | 579.71 (3) |
| Putranjivaceae | 2 | 101.3 | 3.11 | 4.00 | 6.55 | 1.89 | 12.43 | 233.29 (5) |
| Urticaceae | 3 | 153.9 | 1.44 | 6.07 | 3.04 | 2.83 | 11.94 | 31.36 (16) |
| Phyllanthaceae | 6 | 65.7 | 0.83 | 2.59 | 1.75 | 5.66 | 10.00 | 51.49 (14) |
| Anacardiaceae | 6 | 14.6 | 1.53 | 0.58 | 3.23 | 5.66 | 9.47 | 83.11 (9) |
| Sapindaceae | 3 | 43.7 | 2.00 | 1.72 | 4.20 | 2.83 | 8.76 | 195.39 (6) |
| Myrtaceae | 4 | 72.5 | 0.32 | 2.86 | 0.67 | 3.77 | 7.30 | 20.26 (21) |
| Annonaceae | 4 | 51.8 | 0.29 | 2.04 | 0.62 | 3.77 | 6.44 | 13.40 (22) |
| Rutaceae | 6 | 12.9 | 0.02 | 0.51 | 0.04 | 5.66 | 6.21 | 0.844 (29) |
| Myristicaceae | 2 | 39 | 0.87 | 1.54 | 1.84 | 1.89 | 5.26 | 47.57 (15) |
| Acanthaceae | 2 | 69.7 | 0.05 | 2.75 | 0.11 | 1.89 | 4.75 | 1.409 (27) |
| Elaeocarpaceae | 2 | 9.5 | 0.81 | 0.37 | 1.72 | 1.89 | 3.98 | 66.30 (11) |
| Staphyleaceae | 1 | 14.5 | 1.08 | 0.57 | 2.27 | 0.94 | 3.79 | 69.84 (10) |
| Ebenaceae | 2 | 14.5 | 0.61 | 0.57 | 1.28 | 1.89 | 3.74 | 57.38 (13) |

**Table 5.**
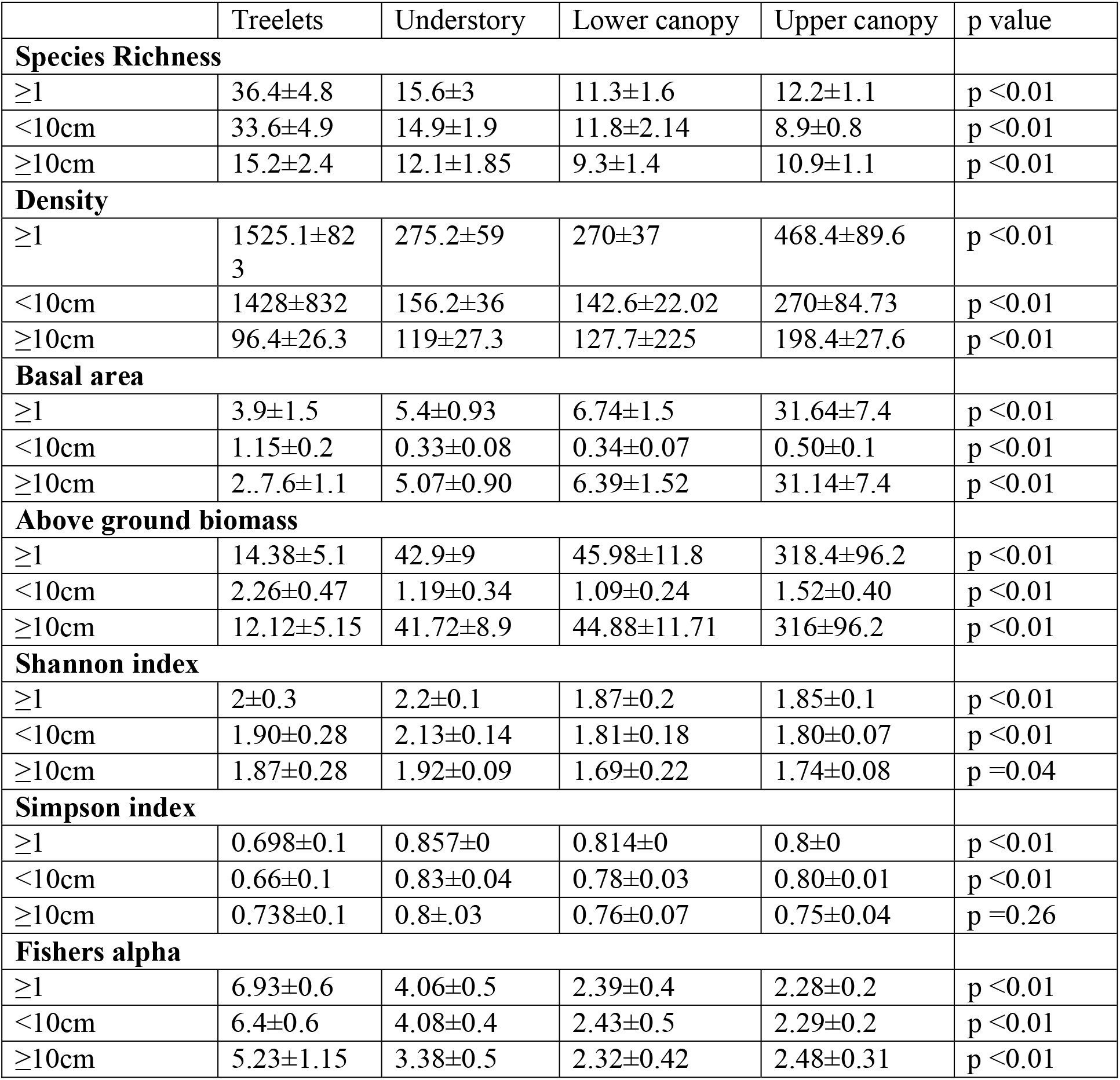
**Distribution of species richness, density, basal area, Above ground biomass, Shannon index, Simpson index, fishers alpha between tree life forms in different diameter class of woody individuals in the Sholayar plot.**

**Table 6:** IVI values of the15 dominant species in seedling, sapling and tree class.

| Species | Seedling | Sapling | Tree |
| --- | --- | --- | --- |
| *Palaquium ellipticum* | 16.79 | 20.5 | 40.01 |
| *Cullenia exarillata* | 20.16 | 22.5 | 33.53 |
| *Mesua ferrea* | 28.57 | 20.5 | 14.41 |
| *Dysoxylum malabaricum* | 21.5 | 6.16 | 5.175 |
| *Vateria indica* | 9.346 | 14.5 | 21.12 |
| *Drypetes venusta* | 6.685 | 7.65 | 16.64 |
| *Syzygium laetum* | 22.2 | 11.2 | 4.967 |
| *Agrostistachys borneensis* | 6.469 | 13.5 | 12.46 |
| *Dimocarpus longan* | 6.377 | 7.74 | 12.24 |
| *Drypetes malabarica* | 10.78 | 6.38 | 10.68 |
| *Aglaia tomentosa* | 13.73 | 23.53 | 8.632 |
| *Antidesma montanum* | 4.085 | 6.75 | 7.607 |
| *Holigarna nigra* | 2.032 | 3.32 | 7.584 |
| *Myristica malabarica* | 1.517 | 12.1 | 7.253 |
| *Turpinia malabarica* | 4.693 | 4.22 | 6.582 |

### Forest structure by life form

Species richness varied significantly among the four life form (p < 0.001). Among the 106 species in the Sholayar 10-ha LTEO plot, there were 54 treelet species, with diameters mostly restricted to <10 cm. These accounted for 60% of the total number of individuals in the plot, but only 8% of basal area and 3.5% of aboveground biomass respectively (Table 2). Seven of the 20 most abundant species in the plot were treelets,

The remaining 52 species comprised 19 understory, 16 lower canopy and 17 upper canopy species, all of which reach more than 10 m in height and dbh**≥**10 cm at maturity. These species represented 40% of the total number of trees, 92% of the total basal area and 96.5% of the total aboveground biomass.

Small-diameter trees included 52 treelets and all upper canopy tree species, as well as 94.73 % and 85.7 % of understory and lower canopy tree species, respectively. In this diameter size class, treelets accounted for 56.27 % individuals, 8.21 % of the basal area, and 3.45 % of the aboveground biomass. Understory and canopy species accounted for only 22.4 % of all trees, 50.64 % of basal area, and 64.22 % of aboveground biomass in these small-diameter trees, whereas accounting for only 25 % of all trees. There were only 41 species of treelets among trees with dbh **≥**10 cm, accounting for 17.80 % individuals and a very small fraction of their total basal area and aboveground biomass in this diameter class. Upper canopy species, on the other hand, accounted for 61.97 percent of the basal area and 64.46 % of the aboveground biomass in this diameter class, accounting for 8.93 % of the individual.

Statistical analysis made by one way ANOVA revealed the means of stem density, basal area and biomass among life form in all three groups (all woody plants and both the diameter class) were statistically significant at P≤0.01. At all diameter classes, values for mean species richness and fishers alpha were much higher than that of the other three life form indicate the more diverse and heterogeneous treelet layer. Furthermore, highest values for Shannon and Simpson diversity index were reported in understory species at both **≥**1 and <10cm dbh classes. Both Shannon and Simpson index among four life forms were not statistically significant at larger diameter class (p= 0.04 and p=0.26 respectively).

### Pattern of regeneration

#### Tree layer

Tree layer is remarked by a total of 59 species and Fisher’s α for this group is 9.426. Based on the IVI value *Palaquium elipticum* was reported dominant species followed by *Cullenia exarillata* and *Vateria indica*. The maximum values of frequency and density were recorded for *Palaquim elipticum* followed by *Cullenia exarillata and Drypetes venusta*. The maximum values of total basal cover were also observed for *Cullenia exarillata* followed by *Palaquim elipticum* and *vateria indica*.

#### Sapling layer

Total 54 species and a Fisher’s α value of 9.05 is reported from sapling layer, the higher value for IVI was reported for *Aglaia tomentosa* followed by *Cullenia exarillata* and *Mesua ferrea*. The IVI value for *Palaquium ellipticum* and *Vateria indica* place them fourth and fifth rank respectively among saplings of all tree species.

#### Seedling layer

In seedling layer contains only 50 species and have Fisher’s α value of 8.71. This layer is remarked with high dominance of *Mesua ferrea* followed by *Syzygium laetum* and *Dysoxylum malabaricum* but the species *Cullenia exarillata and Palaquium ellipticum* have comparatively lower IVI in this class.

## Discussion

Information regarding the floristic composition and diversity of a forest is essential for the ecological research and management. Floristic inventories are prerequisites for constructing the forest models and species distribution patterns. Present study reports the enormous floral diversity of the Sholayar region of southern Western Ghats. According to the family important value index(FIV) that the family Rubiaceae (FIV=44.87) was the most dominant family followed by Sapotaceae (FIV=27.23) and Malvaceae(FIV=25.43). The analysis of the flora of the study area showed that. Rubiaceae was the most represented family followed by Euphorbiaceae and Urticaceae. High densities of Rubiaceae and Euphorbiaceae were also reported from various tropical rain forests (Reynal 1994, ifo 2015).

Although there have been some sound effort in recent past towards long-term monitoring of Western Ghats. Small diameter trees from the evergreen forest of western Ghats received very less attention and they remain under studied. The current study plot is the only one in the Western Ghats’ wet evergreen region where smaller diameter woody plants (gbh **≥** 3.14 cm) were included and completely censused. When compared to other evergreen Asian plots the present study plot has moderate proportion of small diameter woody plants (78.67%) which is above 90% for most of the wet evergreen tropical plots. The basal area and biomass were concentrated in the large diameter class, which affects the ability of the forest to recover from disturbance affecting the large diameter trees. Large diameter trees store bulk of biomass, which is an agreement with previous forest inventories across tropics (Lee et al. 2002). Decreasing the minimum diameter limit to 1 cm, as opposed to the 10 cm limit adopted in most evergreen forest tree surveys, revealed a dense and diverse understory. Advantages of incorporating small diameter woody individuals for sampling forest diversity, dynamics, and demography were revealed by Davies et al. 2021. Similar conclusions can be made from our study. Capturing dynamics in the species diversity is one among the key objectives of long term monitoring of ecosystem. Results from the present study demonstrate that contributions of smaller diameter plants are essential for assessing the diversity of the forest.

The species richness of the present 10 hec permanent plot is relatively good when compared to other large-scale permanent plots (Davies et al. 2021) from southern Western Ghats. Tree density in the sholayar plot was very low when compared to other plots across tropics. For plants with dbh**≥**10 it is comparable with other permanent plots in Western Ghats and has relatively moderate density of woody plants (541 individuals/ha):447 individuals/ha in Varagaliar site and 763 individuals ha^−1^ in Uppangala, central western Ghats India. Around 50% of the species in the plot were treelets that achieve reproductive maturity in the forest under story and rarely attain 10 cm dbh. The small diameter trees also comprised of seedlings and saplings of all other understory and canopy tree species that do regenerate. When considering plant families more than 50% of basal area and biomass were concentrated in four families Malvaceae, Sapotaceae, dipterocarpaceae and calophyllaceae. The treelet species richness of total 54 species in the 10-ha plot reflect a high treelets diversity status of tropicalwet evergreen forest of Western ghats. Fishers α value among life form varies as treelets >under story>lower canopy>upper canopy: this trend is an indication of increasing dominance and decreasing species richness from treelet layer to higher canopy layer. The greater density and species richness in the treelets layer might be the consequences of the high rain fall, soil fertility and higher incidence of endozoochory (Thomas 1999)

Our results showed the Simpson and Shannon diversity index of treelets are lower than under story layer this is because of the higher uniformity of individual distribution among species and higher density. It is evident that at girth class 2 Shannon diversity indices for treelets is higher than that of the other life form this is probably due to increased rarity for treelets in girth class 2. Treelets layer of the plot contribute more to the species richness of both diameter class.

## Regeneration

The regeneration pattern of the species is shown in Table Girth class analysis of the species suggests that dominant species of the tree layer are failed to maintain its dominance in sapling layer and seedling layer. Density of tree species in seedling and sapling layer is crucial because regeneration status of tree species of any forest is determined by recruitment of saplings and seedlings (Singh and Singh 1992, Dharet al. 1997). The species that are currently dominates in tree layer have relatively lower density and dominance in the seedling and sapling class which indicate that, regeneration of these species are affected and in future growth and dominance of these species may effected in the forest. This kind of regeneration dynamics eventually changes the diversity and stand structure of the forest. In seedling layer along with *Mesua ferrea, Syzygium laetum and Dysoxylum malabaricum* dominates in both density and IVI indicates a strong evidence to the regeneration dynamics of the tropical wet evergreen forest. The density values of seedlings and saplings are considered as regeneration potential of the species and presence of good regeneration potential shows suitability of a species to the environment (Saxena and Singh 1984). From the density curve (Figure 2) it is obvious that in tree species density at initial girth class (seedling) is lower than the sapling class indicates an unstable regeneration. Climatic factors and biotic interference influence the regeneration of different species in the vegetation (Saha 2016).

**Figure 2:**
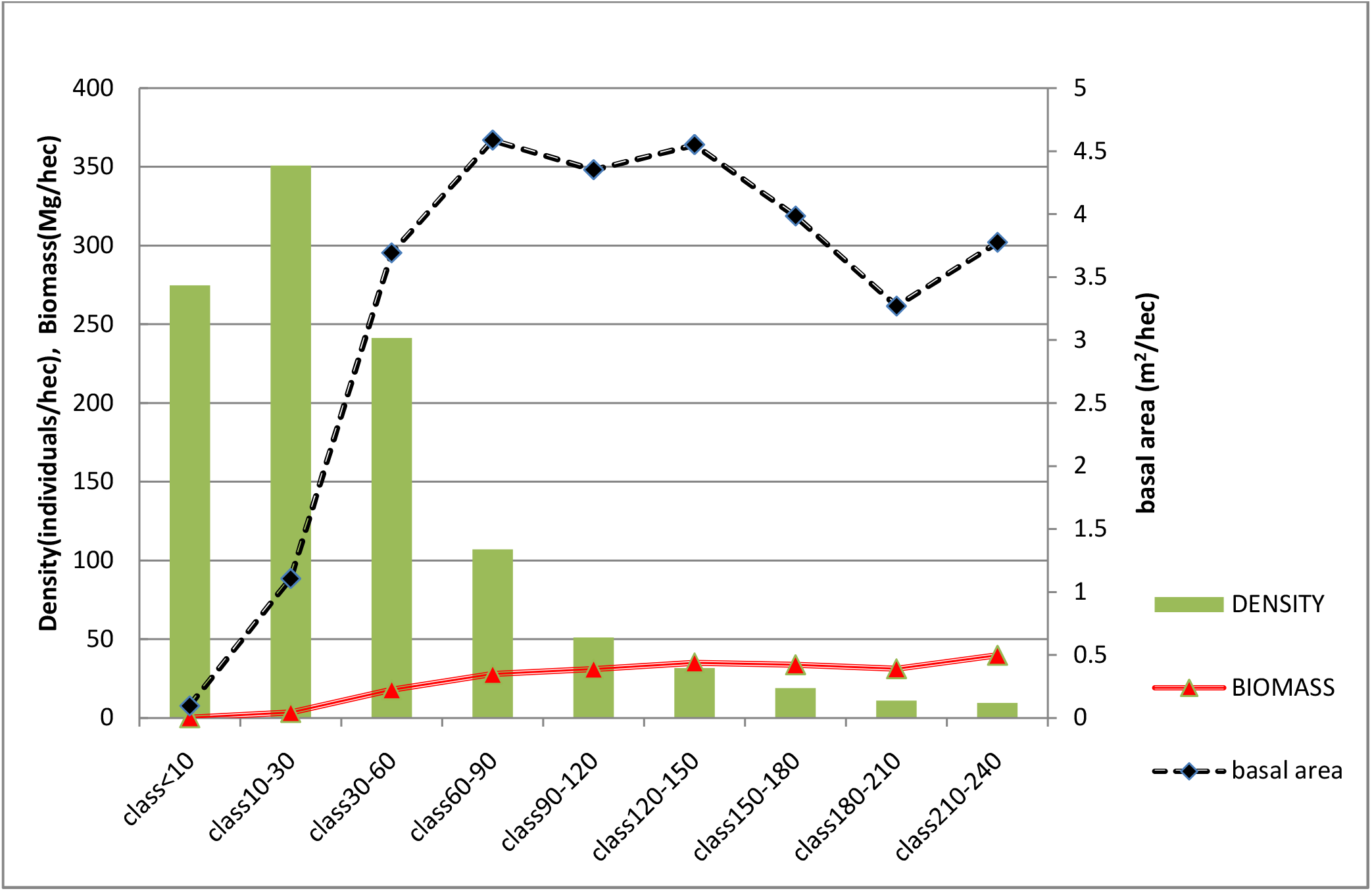
Changes in density, Biomass, basal area across different girth classes.

## Acknowledgement

We thank the Director and former directors, Kerala Forest Research Institute for providing all facilities for the completion of the study. We acknowledge the help of chief wildlife warden, Kerala Forest Department, Govt. of Kerala for field permission and all forest officials including DFO, Vazhachal Forest Division and Forest Range Officer, Sholayar range for their support. We also aknowledge the Kerala Council for Science, Technology, and Environment (KSCSTE).

